# Millisecond-scale detection of mouse ultrasonic vocalizations enables closed-loop experiments

**DOI:** 10.64898/2026.09.24.754020

**Authors:** Ali Mohammadi, Jens Tillmann, Martin K. Schwarz

## Abstract

Mouse ultrasonic vocalizations (USVs) provide a rapidly evolving readout of social interaction but are typically analysed only after acquisition. Here we introduce DeepFisFis, a waveform-based neural network that detects USVs while they are being produced. DeepFisFis classifies consecutive 5 ms audio segments directly from the waveform with high accuracy, processing each segment in approximately 2.5 ms and therefore faster than the incoming audio stream. This enables ongoing vocalizations to guide experimental interventions in real time. In a deployed closed-loop system, detections triggered an external stimulus, demonstrating online control of ongoing vocal behaviour. DeepFisFis also enables event-triggered data acquisition: gating storage around detected calls preserved more than 99% of vocalization time while retaining only approximately 22% of the continuous recording. DeepFisFis thus transforms USVs from a retrospective behavioural readout into a real-time experimental signal for selective acquisition and closed-loop causal interrogation of vocal communication and its underlying neural circuits.

## Introduction

Mice communicate extensively through ultrasonic vocalizations (USVs), calls emitted above the range of human hearing that accompany social behaviours including maternal interaction, courtship and social investigation (Arriaga and Jarvis, 2013; Chabout et al., 2015; Sangiamo et al., 2020). USVs are emitted above about 20 kHz and can extend beyond 110 kHz, and last from a few milliseconds to a few hundred milliseconds, with acoustic structure varying across strain, sex, age and experimental condition (Castellucci et al., 2018; Caruso et al., 2022).

Because USV rate and structure are altered in models of neurological and neurodevelopmental disease, they are widely used as non-invasive behavioural read-outs (Premoli et al., 2021). Reliable use of this read-out depends first on detecting calls accurately, efficiently and at the temporal scale at which they occur.

USV detection remains technically demanding. The high frequencies of mouse vocalizations require high-sampling-rate recordings, generating large volumes of data even in short experiments. Calls can be brief, variable in spectrotemporal shape and produced in rapid sequences, while recordings may also contain ultrasonic artefacts from equipment or from the environment. Manual annotation is therefore slow, difficult to scale and prone to observer-dependent variability (Coffey et al., 2019; Baggi et al., 2023). Automated methods have improved throughput, but their design has largely been optimized for offline analysis of completed recordings (Coffey et al., 2019; Tachibana et al., 2020; Fonseca et al., 2021; Abbasi et al., 2026).

Most established USV detectors operate on time–frequency representations of the signal, typically spectrograms. Some extract handcrafted features, including power in the ultrasonic band, duration, contour or harmonic structure, and apply user-defined thresholds or image-processing steps (Holy and Guo, 2005; Tachibana et al., 2020; Ashley et al., 2021; Pessoa et al., 2022). Others apply deep learning, either to detect calls directly from spectrogram images (Coffey et al., 2019) or to reject noise and classify candidates found by image processing (Fonseca et al., 2021). These approaches have enabled increasingly accurate offline analysis, but they rely on computing an intermediate representation and are generally applied after acquisition is complete. This limits their use in closed-loop experiments, where the experimental setup must detect an ongoing vocalization with sufficiently low latency to trigger a predefined response.

Systems capable of online detection have been described, with throughput sufficient to keep pace with acquisition (Steinfath et al., 2021; Stoumpou et al., 2023). Their reported analysis windows are nonetheless long relative to the duration of a mouse USV. For experiments in which a stimulus must be associated with the call that elicited it, what matters is not only that audio can be processed as fast as it is acquired, but how soon after a call begins a decision becomes available.

We address this need with DeepFisFis, a waveform-domain detector designed for real-time USV detection. DeepFisFis divides the incoming audio stream into short segments and uses a one-dimensional convolutional neural network (1D-CNN) to estimate the probability that each segment contains part of a USV. Suprathreshold segments are then temporally post-processed and merged into discrete detected calls. By learning directly from the audio waveform, the model avoids hand-tuned feature extraction and predefined acoustic measurements. Bypassing the time–frequency transform also removes its minimum window length, allowing shorter analysis segments, and keeps per-segment processing small enough for low-latency operation (Steinfath et al., 2021).

Here we introduce DeepFisFis, a compact 1D-CNN that detects mouse USVs directly from 5 ms waveform segments and is designed to act during rather than after acquisition. We first establish that direct waveform analysis accurately recovers USVs and that each 5 ms segment can be processed faster than it is acquired. We then demonstrate this capability in a deployed closed-loop configuration in which detected USVs trigger an acoustic stimulus, and in event-triggered acquisition that retains call-centred data while substantially reducing storage.

Finally, we show that the waveform-based detector can be adapted across sampling rates and acquisition settings without additional recordings. DeepFisFis therefore turns ultrasonic vocalizations from a retrospective behavioural readout into a real-time experimental signal, providing a basis for closed-loop causal experiments and selective multimodal acquisition during social interaction.

## Results

### Direct waveform analysis detects USVs on a 5 ms timescale

We developed DeepFisFis as a one-dimensional convolutional neural network (1D-CNN) for detecting mouse ultrasonic vocalizations directly from the time-domain audio waveform (Figure 1A). Rather than first converting the audio into a spectrogram or other derived representation, DeepFisFis operates directly on the waveform. Audio is divided into 5-ms segments, each of which is independently classified according to whether it overlaps a USV. Consecutive positively classified segments are subsequently merged to reconstruct individual vocalization events.

**Figure 1.**
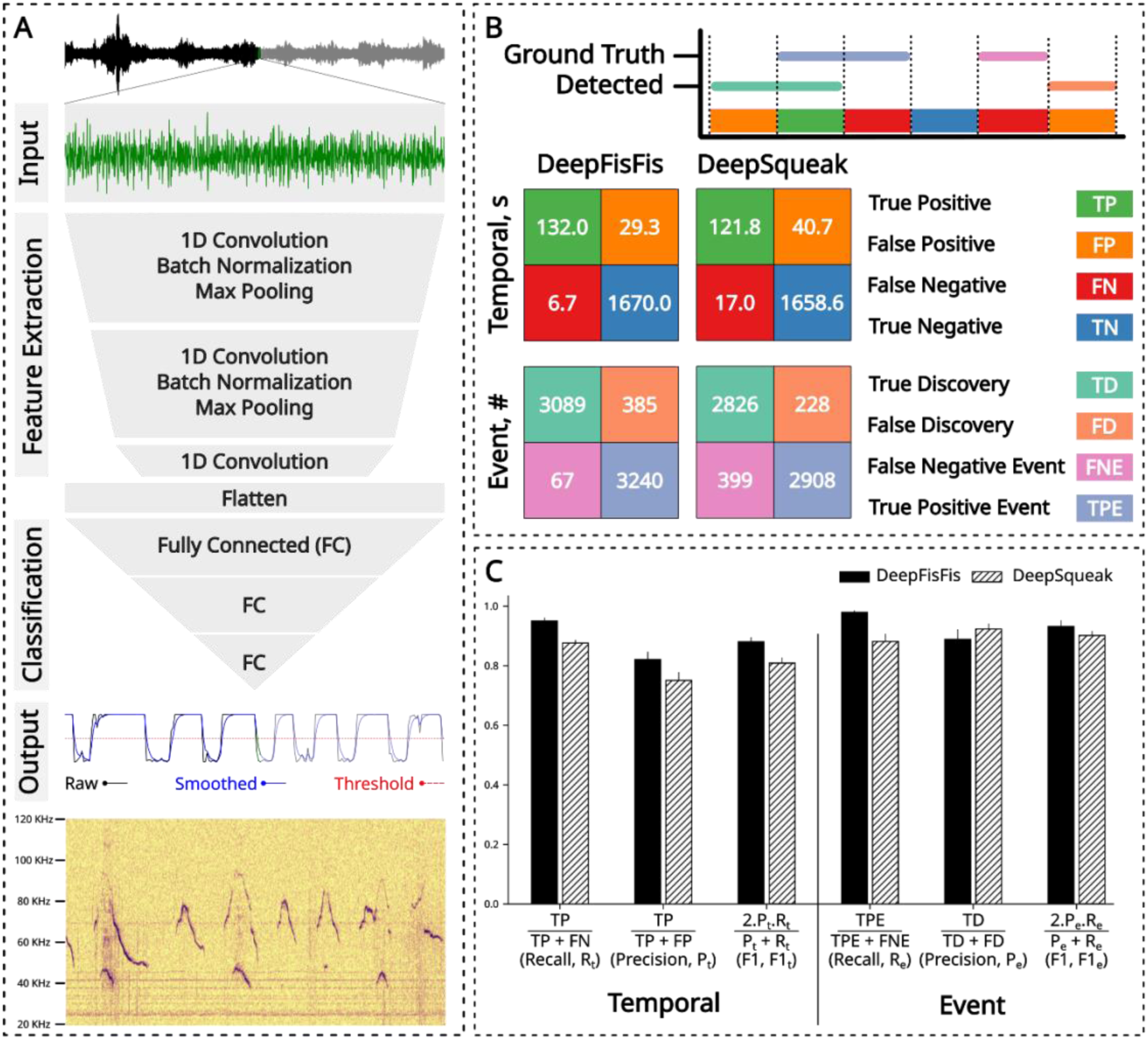
Network architecture and detection accuracy at 250 kHz. **(A)** Schematic of the 1D-CNN at the core of the DeepFisFis pipeline. The network classifies 5-ms waveform segments, outputting the probability that each overlaps a USV. Consecutive probabilities are smoothed by a causal exponential filter and thresholded at 0.5; positive segments separated by 5 ms or less are merged into vocalization events. The lower panel shows the temporal alignment of model outputs with the corresponding spectrogram. **(B)** Temporal and event-based confusion matrices for DeepFisFis and DeepSqueak on annotated recordings from multiple sources (scoring schemes defined in Materials and methods, ‘Evaluation and comparison’). Matrix entries are pooled across folds. **(C)** Recall, precision, and F1 score derived from the confusion matrices in (B), shown as mean ± SD across five folds.

This architecture was designed to combine accurate USV detection with continuous online processing. Because each 5-ms segment can be classified as soon as it is acquired, DeepFisFis does not require a complete vocalization or an extended audio interval to be recorded before analysis begins. The segment-based architecture therefore operates on a timescale comparable to individual USV production, making real-time detection feasible.

We first asked whether direct classification of the audio waveform was sufficient to achieve detection performance comparable to an established USV detection tool, DeepSqueak (Coffey et al., 2019). To this end DeepFisFis was trained and evaluated on 250-kHz recordings using 5-fold cross-validation and benchmarked against DeepSqueak on the same recordings.

Performance was quantified using complementary temporal and event-based metrics, assessing both agreement with the annotated audio signal and recovery of individual vocalization events (Figure 1B,C).

DeepFisFis achieved a temporal F1 score of 0.88 ± 0.01 (precision 0.82, recall 0.95). At the event level, DeepFisFis reached an F1 score of 0.93 ± 0.02 (precision 0.89, recall 0.98; Figure 1B,C). DeepSqueak achieved a temporal F1 score of 0.81 ± 0.02 (precision 0.75, recall 0.88) and an event-based F1 score of 0.90 ± 0.01 (precision 0.92, recall 0.88; Figure 1B,C). Note that all values are mean ± SD across folds.

Thus, direct analysis of short waveform segments was sufficient to recover USVs with high temporal and event-level accuracy. In particular, the event-level recall of 0.98 indicated that nearly all annotated vocalization events were detected. These results establish that transforming the input audio into an intermediate form is not required for accurate USV detection and that a 1D-CNN operating directly on the audio waveform can achieve performance comparable to or exceeding that of DeepSqueak.

### Five-millisecond waveform analysis runs faster than audio acquisition

For subsequent real-time experiments, we adapted DeepFisFis to 384-kHz recordings using the transfer procedure described below. The short input window used by DeepFisFis was designed to permit continuous processing during data acquisition. To verify this, we measured the time required to classify individual audio segments and compared the deployed implementations of DeepFisFis and DeepSqueak across different input durations (Figure 2).

**Figure 2.**
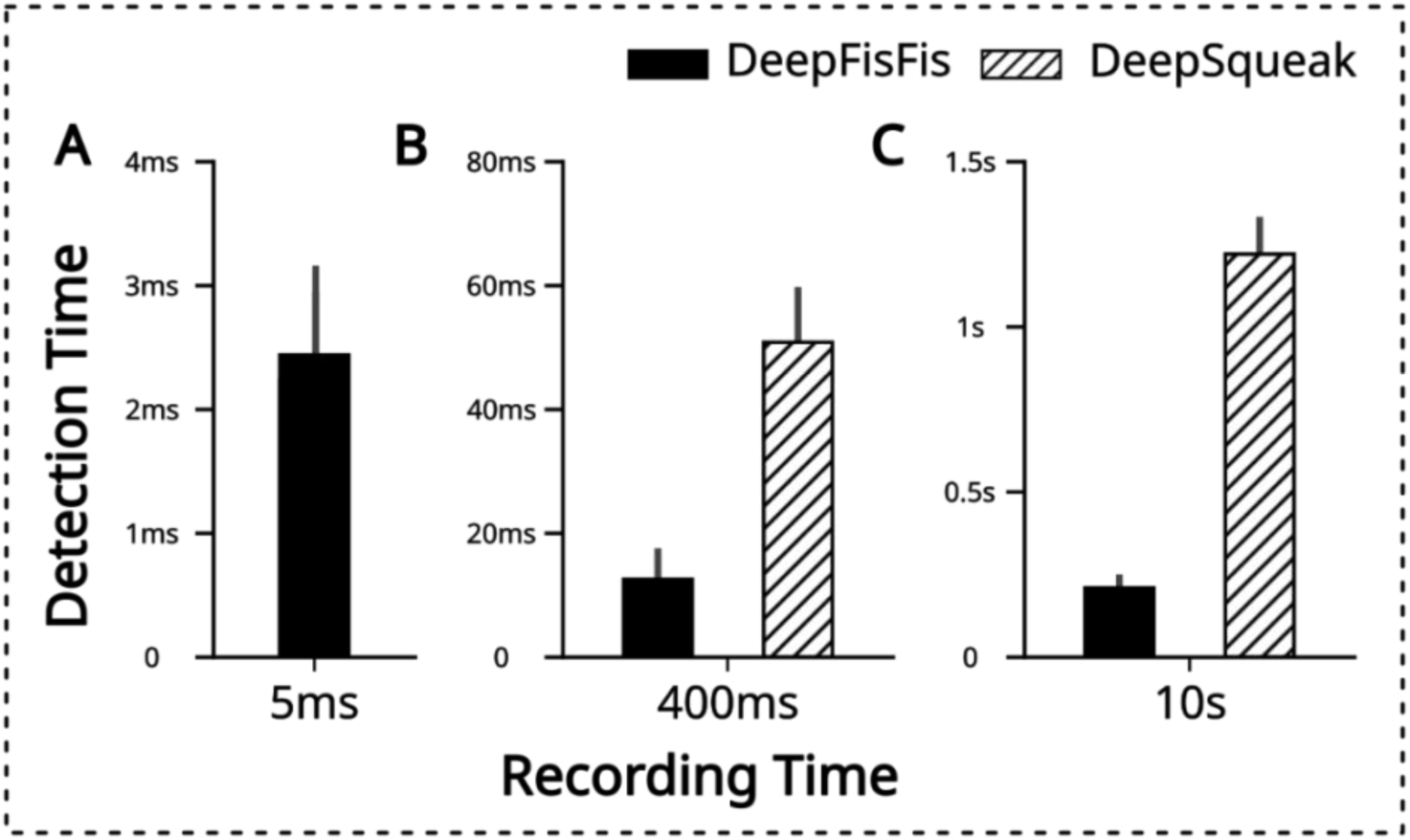
Processing speed comparison, DeepFisFis vs. DeepSqueak. (**A–C**) Processing time per input length for DeepFisFis at input lengths of 5 ms (A), 400 ms (B), and 10 s (C), measured using the 384-kHz variant on 384-kHz recordings. DeepSqueak was compared at 400 ms and 10 s, as it was not applicable at 5 ms. Bars show mean processing time averaged over 1000 repetitions; error bars show SD.

To provide a conservative benchmark, measurements were performed using the 384-kHz variant of DeepFisFis, which must process longer input segments sample-wise and employs a slightly larger model than the 250-kHz version.

DeepFisFis classified each 5-ms waveform segment in approximately 2.5 ms on a single GPU (Figure 2A). Processing was therefore completed substantially faster than the time required to acquire the corresponding audio segment. This allowed successive waveform segments to be continuously analyzed without accumulating an increasing processing delay during recording. When processing longer recordings, DeepFisFis also processed audio substantially faster than DeepSqueak. For the shortest input duration accepted by DeepSqueak, 400 ms, DeepFisFis was approximately four times faster (Figure 2B). At an input duration of 10 s, this difference increased to approximately five-fold (Figure 2C). These measurements reflect the standard working implementations of both approaches rather than a controlled comparison of theoretical computational complexity. Nevertheless, the difference in processing time was large and consistent across all tested input durations.

### Real-time USV detection drives closed-loop experimental responses

Together, these results demonstrate that DeepFisFis processes audio faster than it is acquired, as required for real-time operation. To test this in a deployed closed-loop system, we implemented a setup in which pre-recorded USVs were played back and detected in real time by DeepFisFis, triggering a tone upon detection. We additionally characterised the hardware and software delays contributing to overall latency independently of the detection algorithm, here referred to as system latency. System latency was estimated in a similar experimental paradigm in which a tone was triggered upon arrival of the first audio chunk at the detection system, bypassing DeepFisFis entirely (Figure 3A). Across 1000 repetitions, system latency was 128 ± 44 ms (mean ± SD). When DeepFisFis was included, the detection pipeline added only a small additional delay (Figure 3B), confirming that system latency is the primary contributor to overall end-to-end delay. Closed-loop timing was therefore set by the acquisition and output hardware rather than by the detector, and must be improved independently of the detection method.

**Figure 3.**
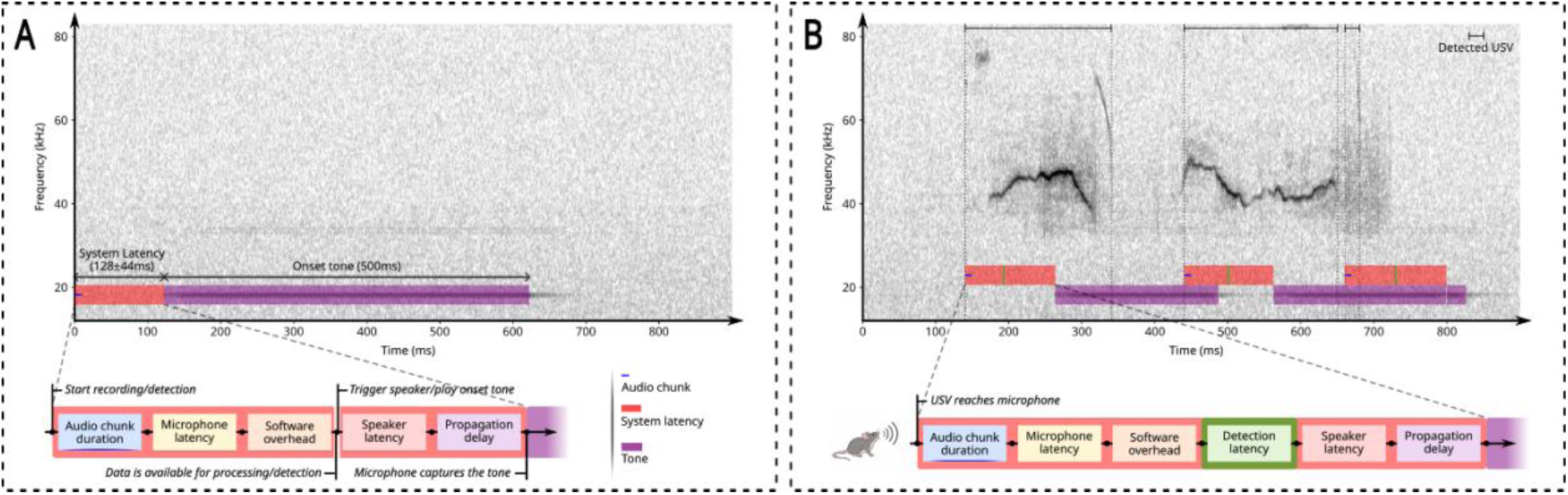
DeepFisFis enables closed-loop USV detection. Latency was characterised in a playback setup in which audio was acquired continuously and a detection triggered a tone through a separate speaker, with both the played-back vocalizations and the tone captured by the same microphone. **(A)** System latency, comprising hardware and software delays independent of the detection algorithm, was estimated by measuring the delay between recording onset and detection of a calibration/onset tone (128 ± 44 ms, mean ± SD, n = 1000 repetitions). The schematic shows the contributing temporal components. **(B)** In the full closed-loop setup, pre-recorded USVs were detected in real time by DeepFisFis and used to trigger a speaker tone. The schematic shows the additional detection latency introduced by DeepFisFis alongside the system latency components from (A). In both panels the upper trace is the spectrogram of one example recording, with the latency components drawn below.

### Real-time detection enables selective event-triggered data acquisition

Continuous high-sampling-rate recordings generate large data volumes, although only a relatively small fraction of the recording may contain vocalizations. By processing the audio stream in real time, DeepFisFis can serve as an event trigger, initiating data storage only when a USV is detected, together with a configurable buffer before and after each event.

In this mode, detection of a USV triggers storage of the corresponding audio interval to be written to disk together with a configurable temporal buffer preceding and following the detected event (Figure 4A). Intervals without detected vocalizations are not retained, effectively converting continuous audio acquisition into an event-triggered recording scheme while preserving temporal context around each detected vocalization.

**Figure 4.**
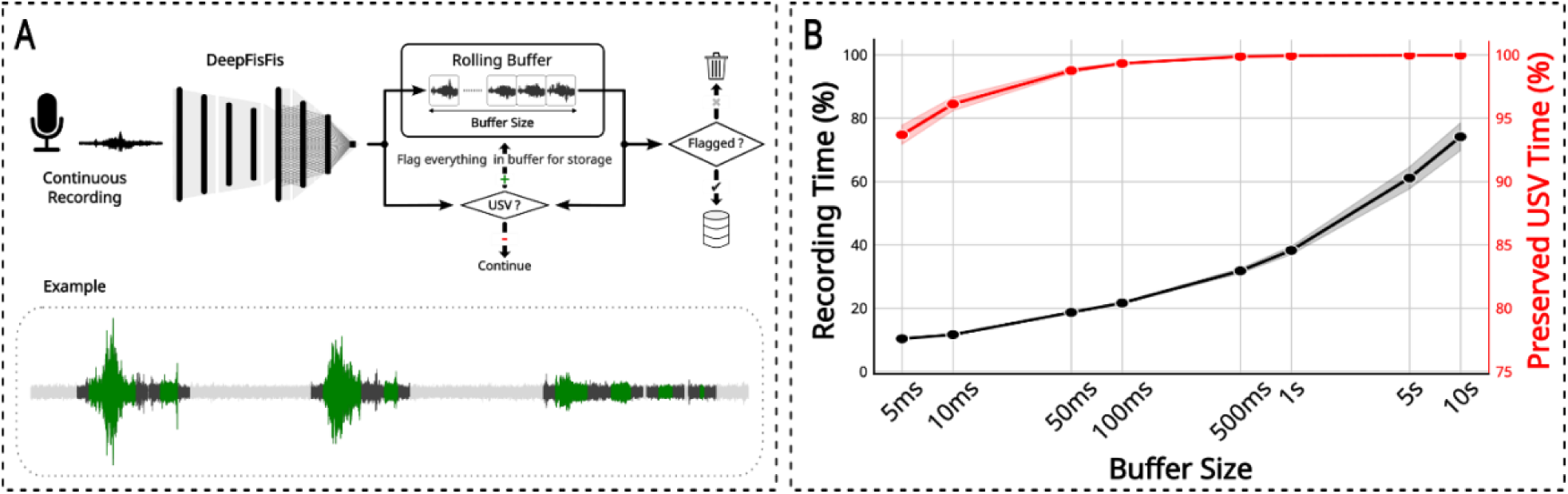
Selective recording using real-time USV detection. **(A)** Incoming audio is continuously processed by DeepFisFis and held in a rolling buffer of configurable size. When a USV is detected, the buffered audio preceding the event and a buffer of the same size following it are flagged for storage. The lower panel illustrates an example in which USV-containing intervals (green) are retained and silent periods (grey) are discarded. **(B)** Recording time retained (black, left axis) and USV time preserved (red, right axis) as a function of buffer size, measured using the 384-kHz variant on 384-kHz recordings. Shaded areas indicate SD across folds.

The fraction of the original recording retained depends on the duration of the selected temporal buffer (Figure 4B). With a short 5-ms buffer, approximately 10% of the original recording duration would be stored while preserving approximately 94% of annotated USV time. Increasing the buffer progressively increased the retained fraction at the cost of additional storage, reaching approximately 22% retention while preserving over 99% of USV time at a 100-ms buffer.

These results demonstrate that DeepFisFis can use ongoing vocalization detection to control data acquisition in real time. Event-triggered recording substantially reduces the volume of stored audio and any simultaneously acquired modalities while retaining vocalization-containing intervals and their surrounding temporal context.

### Waveform-based detection transfers across sampling rates without new recordings

USV recordings are acquired using different sampling rates, microphones, and acoustic environments, raising the question of whether the DeepFisFis model trained under one acquisition condition can be transferred to another without complete retraining. We therefore tested whether the model trained on 250-kHz recordings could be adapted to process recordings acquired at 384 kHz.

The 384-kHz recordings were not included during initial model training. Instead, the pretrained 250-kHz model was adapted using the transfer-learning procedure described in Materials and methods (‘Model fine-tuning (384 kHz)’), without acquiring additional recordings. Fine-tuning used only the 250 kHz training recordings resampled to 384 kHz, so every 384 kHz recording remained unseen by the model and served as an independent test set. The adapted model was subsequently evaluated on held-out 384-kHz recordings, most of which were acquired in our own laboratory rather than from the sources used for training, using the same temporal and event-based metrics employed for the 250-kHz dataset (Figure 5A,B).

**Figure 5.**
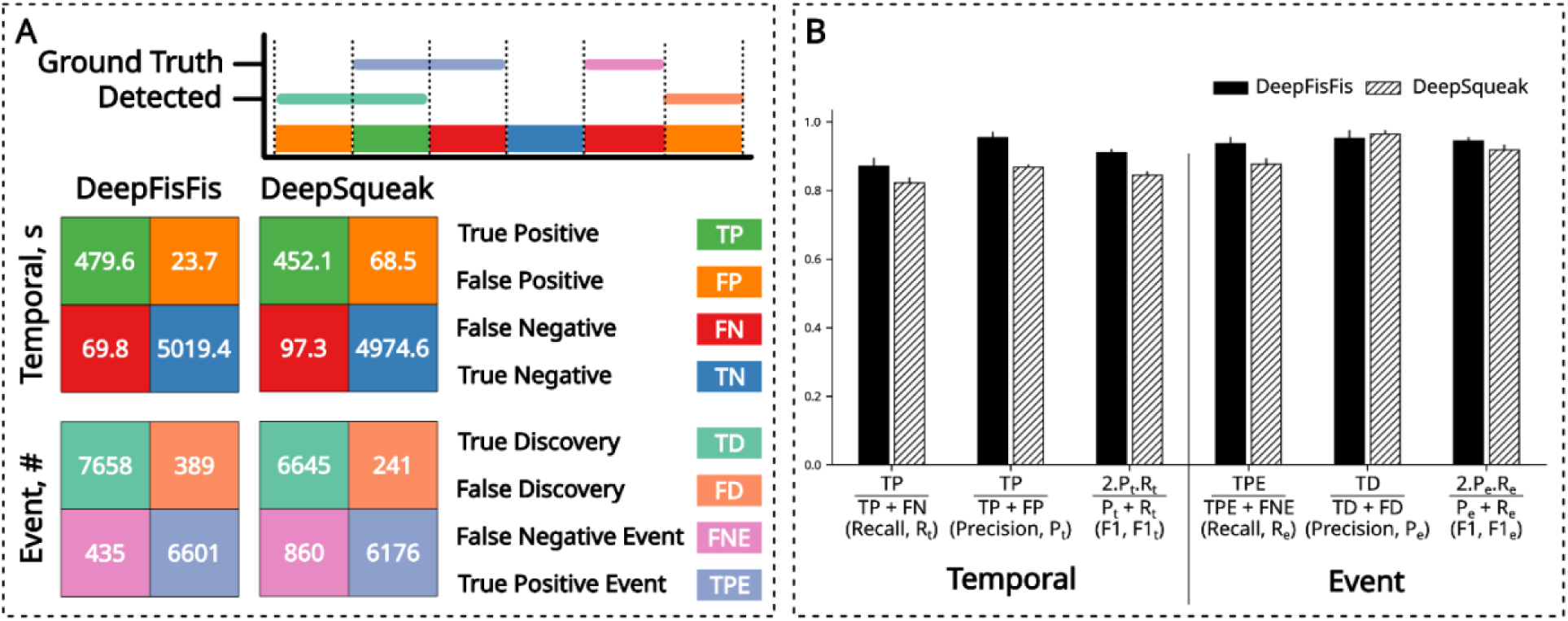
Detection accuracy after transfer to 384 kHz. **(A)** Temporal and event-based confusion matrices for DeepFisFis and DeepSqueak on held-out 384-kHz annotated recordings (scoring schemes defined in Materials and methods, ‘Evaluation and comparison’). Matrix entries are pooled across folds. **(B)** Recall, precision, and F1 score derived from the confusion matrices in (A), shown as mean ± SD across five folds.

Following transfer, DeepFisFis achieved a temporal F1 score of 0.91 ± 0.01 (precision 0.95, recall 0.87) and event-based performance reached an F1 score of 0.94 ± 0.01 (precision 0.95, recall 0.94; Figure 5A,B). On the same recordings, DeepSqueak achieved a temporal F1 score of 0.84 ± 0.01 (precision 0.87, recall 0.82) and an event-based F1 score of 0.92 ± 0.01 (precision 0.96, recall 0.88; Figure 5A,B).

Importantly, adapting DeepFisFis to the higher sampling rate did not reduce detection performance relative to the original 250-kHz recordings. Temporal and event-based F1 scores remained at least as high after transfer. Thus, a model initially trained on 250-kHz audio could be adapted to process 384-kHz recordings by fine-tuning on the original training material resampled to the new rate, while maintaining high USV detection accuracy.

Together, these results show that direct waveform analysis with a segment-based 1D-CNN enables accurate USV detection without engineered features or intermediate representations. Processing is sufficiently fast to detect vocalizations as they are produced, enabling closed-loop triggering and event-triggered acquisition, while the same waveform representation can be adapted across acquisition conditions and sampling rates with only limited fine-tuning.

## Discussion

DeepFisFis shifts mouse USV analysis from retrospective detection to real-time experimental control. By classifying 5 ms waveform segments directly, the system detects ongoing vocalizations faster than the corresponding audio is acquired, allowing the acoustic signal itself to trigger experimental responses or determine which data are retained. We show that this low-latency design maintains high detection accuracy on manually annotated recordings and adds only a small delay to a deployed closed-loop system, in which hardware rather than inference sets the dominant latency. Applied to event-triggered recording, the same real-time detections preserve nearly all vocalization content while retaining only a fraction of the continuous audio. Finally, the detector transfers from 250 to 384 kHz without acquiring new training data, indicating that the waveform-based representation can be adapted across acquisition settings. Together, these findings establish DeepFisFis not simply as a faster USV detector, but as an acquisition-stage tool that makes vocalizations available as real-time experimental events.

The main contribution of DeepFisFis is therefore not that it replaces established USV analysis pipelines (Coffey et al., 2019; Tachibana et al., 2020; Fonseca et al., 2021; Stoumpou et al., 2023), but that it addresses a different experimental requirement. Tools such as DeepSqueak provide rich offline analysis workflows, including user interfaces and downstream call analysis. DeepFisFis is instead optimized for low-latency detection from a continuous input stream. This distinction is important for closed-loop experiments, in which detection must occur during the vocalization rather than after completion of the recording. In the present implementation a decision requires only a single 5 ms segment containing at least 2.5 ms of vocalization, the criterion used to label training segments, so a detection becomes available 5–7.5 ms after call onset, depending on where the call begins within the segment. This is shorter than all but the briefest mouse USVs, which extend to a few hundred milliseconds (Castellucci et al., 2018). In a deployed system the total delay additionally includes the latency of the acquisition and actuation hardware. In our configuration this amounted to 128 ± 44 ms (Figure 3). Whether feedback can be delivered within a single call therefore depends on reducing system latency, for example with dedicated low-latency acquisition hardware (Durieux et al., 2026), rather than on accelerating the detector further.

Online detection has been demonstrated before, but at temporal resolutions set by the analysis window rather than by the vocalization. AMVOC processes audio in 750 ms blocks (Stoumpou et al., 2023), and the DAS configuration reported for mouse USVs analyses 8,192-sample chunks, 27.3 ms at 300 kHz, with 7 to 15 ms of inference per chunk (Steinfath et al., 2021). Mouse USVs are brief and densely packed: inter-syllable intervals peak near 20 ms, and intervals beyond 160 ms are conventionally taken to terminate a sequence (Hertz et al., 2020). A detection delivered tens or hundreds of milliseconds after onset therefore arrives during a later syllable, or after the sequence has ended. DeepFisFis reaches a decision within a single segment, 5 to 7.5 ms after onset. The contribution is therefore a detection whose delay is set by the onset of the call rather than by the duration of the analysis window.

A second advantage follows from the waveform-based design. Because the detector operates directly on minimally preprocessed audio, it avoids the computational cost of generating spectrograms and reduces dependence on hand-tuned parameters (Steinfath et al., 2021). The features used for detection are learned internally by the network, which makes the approach adaptable. In concurrent work, a one-dimensional convolutional network applied to consecutive waveform chunks has been used to segment birdsong syllables online and trigger acoustic feedback during learning experiments (Riekers et al., 2026). While applied to a different species and a longer timescale, that work arrives at the same design choice, which suggests that frame-wise classification of the waveform is a general basis for closed-loop acoustic experiments rather than a solution specific to ultrasonic vocalizations.

The segment length places a further bound on what the network can learn. A mouse USV typically lasts tens to hundreds of milliseconds, so a 5 ms segment contains only a small fraction of a call and cannot represent its frequency trajectory. What the classifier can learn from such a segment is therefore local waveform structure, the tonal, narrowband character shared by vocalizations, rather than the morphology of any particular call type. Temporal context enters only through the sequence of segment probabilities, in which a positive segment raises the evidence for those adjacent to it, so no spectrotemporal feature extending beyond a single segment is represented. This is consistent with the higher recall and slightly lower precision observed relative to DeepSqueak, although the balance between the two also depends on the detection threshold. The 384 kHz evaluation points in the same direction: accuracy was maintained across recording conditions, and without new annotated calls.

A detector that has learned the morphology of the call types present in its training data is, by construction, best suited to recognizing those types, and may be less reliable for calls whose form departs from them. Robustness to such calls was therefore a design consideration here, since a representation built from local waveform structure rather than from call morphology should be less tied to the repertoire it was trained on. Such departures may arise in contexts that have not been systematically studied, in disease models where altered vocal output is itself the phenotype under investigation, or in social settings where two or more animals vocalize at the same time. Whether this robustness is realized in such settings remains to be established and is a target for future work.

The same representation may also provide a useful starting point for related tasks such as call-type classification or emitter-specific analysis, which at present relies on microphone arrays and offline beamforming (Sangiamo et al., 2020; Sterling et al., 2023). This is consistent with the broader project goal of extending real-time USV detection toward classification, localization and integration with markerless behavioural tracking during social interaction.

However, apart from the closed-loop and event-triggered recording modes demonstrated here, these extensions remain future developments rather than demonstrated functions of the current detector.

Several limitations qualify the present results. First, the comparison with DeepSqueak is practical rather than fully controlled. The tools differ in implementation, training history, and intended use. DeepSqueak was not designed for very short streaming inputs, whereas DeepFisFis was optimized for that setting. The benchmark therefore supports the conclusion that DeepFisFis achieves comparable accuracy and substantially faster processing in the tested deployment, but it should not be interpreted as an isolated comparison of algorithmic complexity. To our knowledge, online detection of mouse USVs has been reported by two other methods (Steinfath et al., 2021; Stoumpou et al., 2023), and we are not aware of either having been applied in a closed-loop experiment. Neither was run here, and the comparisons drawn with them in the Discussion rest on the configurations reported in their publications rather than on measurements made under matched conditions. Second, the validation dataset was limited. We here used manually annotated 250 kHz recordings and cross-validation, but broader validation across laboratories, recording devices, strains, behavioural contexts and noise conditions will be needed to establish general robustness. Folds were constructed by dividing concatenated recordings into consecutive 10 s parts, so parts recorded from the same animals in the same acoustic environment can fall into different folds. The 250 kHz estimates should therefore be read as within-dataset performance. This is complemented by the 384 kHz evaluation, in which the model was fine-tuned only on the resampled 250 kHz training material and then applied to recordings it had never seen, including recordings acquired in a different laboratory with different animals and different equipment. Accuracy did not decrease under these conditions, which suggests generalization to new acquisition configurations, such as a different acoustic environment or sampling rate. Third, the 384 kHz evaluation rests on three recordings, and broader testing across acquisition settings would strengthen it. Fourth, robustness to calls whose form departs from those represented in the training data was not evaluated, and cannot be evaluated against these data: the ground truth consists of manual annotations, which by construction cannot include a detection that a human annotator would not have made. Finally, onsets and offsets are quantized at the 5 ms segment resolution, and event matching is many-to-many, e.g. at 250 kHz, 3,240 of 3,307 annotated calls were recovered by 3,089 of 3,474 predicted calls, indicating that closely spaced calls were sometimes merged into a single detected event.

The event-triggered recording analysis illustrates a direct practical consequence of real-time detection. High-sampling-rate USV recordings can become large in long behavioral or multimodal experiments, especially when synchronized with video, electrophysiology, or imaging. By detecting calls online and retaining only call-centred intervals with configurable temporal buffers, DeepFisFis can reduce storage demands before an offline processing step is required. This use case does not yet constitute a full closed-loop behavioral experiment, but it demonstrates that the detector can already act during acquisition and can support experimental designs that depend on online acoustic monitoring.

In conclusion, DeepFisFis shows that accurate mouse USV detection can be performed directly on the waveform, with a decision available within a single 5 ms segment of a call’s onset. It performs comparably to an established offline detector on manually annotated recordings, and adapts to new sampling rates and recording conditions without acquiring new training data. In a deployed closed loop the remaining delay was set by the acquisition and actuation hardware rather than by detection, which places the limit on closed-loop USV experiments in the instrumentation rather than in the analysis. The next step is to test the method in live closed-loop experiments with freely behaving animals and to extend the learned representation toward real-time call classification, emitter localization and integration with behavioural tracking. Together, these developments would turn DeepFisFis from a fast detector into a platform for interactive studies of vocal communication during social behavior.

## Materials and methods

### Detection pipeline

DeepFisFis processes audio in a four-stage pipeline: segmentation, preprocessing, classification, and merging (Figure 1A). It operates directly on the raw waveform without spectral transformation. The ordering of segmentation and preprocessing, and the role of the merging stage, may vary with deployment context.

#### Segmentation

The recording is divided into consecutive, non-overlapping 5 ms segments (1,250 samples at 250 kHz and 1,920 at 384 kHz). Short segments limit per-segment computational cost and cap the latency before a segment can be classified.

#### Preprocessing

Each segment is bandpass filtered to suppress out-of-band energy (e.g. cage and handling noise) using an eighth-order zero-phase Butterworth filter applied with forward-backward filtering (filtfilt). We used cutoffs of 15–124 kHz for 250 kHz recordings and 15– 170 kHz for 384 kHz recordings. Filtering is applied independently to each segment, so the operation is non-causal only within the 5 ms segment already held in memory and does not introduce additional latency. Each filtered segment is then scaled to unit Euclidean norm so that the classifier responds to waveform shape rather than absolute level, which varies across subjects and recording conditions.

#### Classification

The 1D-CNN (‘Network architecture and training’, below) assigns each segment a probability that it overlaps a USV call. Consecutive probabilities form a time series, which is smoothed by a causal exponential filter (α = 0.5) to suppress isolated fluctuations. A segment is then labelled positive when its smoothed probability exceeds the decision threshold (0.5). The threshold can be raised to reduce false positives or lowered to reduce false negatives, allowing the experimenter to tune sensitivity for the application.

#### Merging

In real-time applications, segments classified as positive can directly trigger an external event (e.g., in closed-loop experiments). However, when call boundaries are required, consecutive positive segments are merged into a single detected call defined by its onset and offset. Gaps of 5 ms or less between detections (corresponding to one negative segment) are bridged, provided the gap has not yet been committed to output.

### Network architecture and training

This section describes the architecture and training of the 1D-CNN used in the classification stage of the DeepFisFis pipeline. The network is first trained on 250 kHz recordings and subsequently adapted to 384 kHz via transfer learning.

#### Network architecture

The classifier is a feedforward 1D-CNN adapted from an architecture for environmental sound classification (Abdoli et al., 2019; Figure 1A). It comprises two blocks: a feature extraction block of three convolutional layers with rectified-linear activations, the first two of which are followed by batch normalisation (Ioffe and Szegedy, 2015) and max-pooling; and a classification block of two fully connected layers with dropout (Srivastava et al., 2014), followed by a sigmoid output unit. The detailed layer configuration and parameter counts for the two model variants (250 and 384 kHz) are given in Table 1.

**Table 1.** Architecture of the DeepFisFis 1D-CNN. *k* and *s* denote kernel size and stride, respectively. The Parameters column reports the total count per layer, including non-trainable batch normalisation statistics. Dual values (e.g. 528 / 1,472) indicate layers that differ between the two model variants (250 and 384 kHz).

| Block | Layer | Configuration | Output shape | Parameters |
| --- | --- | --- | --- | --- |
| Input | — | 1,250 / 1,920 samples | (1,250, 1) / (1,920, 1) | — |
| Feature extraction | Conv1D + ReLU | 16 filters, $k = 32$, $s = 2$ / $k = 91$, $s = 3$ | (610, 16) | 528 / 1,472 |
|  | BatchNorm | — | (610, 16) | 64 |
| | MaxPool + ReLU | pool = 2, $s = 2$ | (305, 16) | — |
| | Conv1D + ReLU | 32 filters, $k = 16$, $s = 2$ | (145, 32) | 8,224 |
|  | BatchNorm | — | (145, 32) | 128 |
| | MaxPool + ReLU | pool = 2, $s = 2$ | (72, 32) | — |
| | Conv1D + ReLU | 64 filters, $k = 8$, $s = 2$ | (33, 64) | 16,448 |
|  | Flatten | — | (2,112) | — |
| Classification | Dense + ReLU | 128 units | (128) | 270,464 |
|  | Dropout | rate = 0.25 | — | — |
|  | Dense + ReLU | 64 units | (64) | 8,256 |
|  | Dropout | rate = 0.25 | — | — |
|  | Dense + Sigmoid | 1 unit | (1) | 65 |
| Total |  |  |  | 304,177 / 305,121 |
| Trainable |  |  |  | 304,081 / 305,025 |

#### Model training (250 kHz)

The model was trained on annotated recordings acquired at 250 kHz. Recordings were bandpass-filtered as described in ‘Detection pipeline’, segmented into 5 ms training segments, and amplitude-normalised. Segments were labelled as non-USV if they had no temporal overlap with any annotated call, and as USV if they overlapped a call by more than 50% of the segment duration. Segments with intermediate overlap were discarded.

Because USVs are sparse and transient, non-USV segments constitute the majority class. Class imbalance was addressed by undersampling the majority (non-USV) class to a 1:1 ratio. A validation set of equivalent class composition was constructed from separate audio files and used for early stopping during training.

The network was trained to minimise binary cross-entropy using the Adam optimiser (Kingma and Ba, 2015) with a learning rate of 10^−4^ and a batch size of 100. Training ran for up to 100 epochs with data shuffled at each epoch. Early stopping monitored the area under the ROC curve (AUC) on the validation set, halting if no improvement was observed for five consecutive epochs and restoring the weights from the best-performing epoch.

#### Model fine-tuning (384 kHz)

The 384 kHz variant was initialised from the trained 250 kHz model. The same annotated 250 kHz dataset resampled to 384 kHz was used, without acquiring new data. The model architecture was adapted for the longer input by scaling the stride of the first convolutional layer with the sampling-rate ratio and enlarging the kernel size to preserve the output length. The corresponding weights were resampled to the new temporal resolution and padded to the target kernel size. All remaining layers retained their weights from the 250 kHz model unchanged. This approach tests whether the representation learned at 250 kHz transfers to a higher sampling rate without additional data collection.

Fine-tuning proceeded in two steps. In the first step, only the first convolutional layer was trainable while all remaining layers were frozen. The model was trained for five fixed epochs using the Adam optimiser (learning rate 5 × 10^−5^) and binary cross-entropy loss. In the second step, all layers of the feature extraction block were made trainable while the classification block was frozen. Training followed the same procedure as the 250 kHz model: Adam optimiser (learning rate 10^−6^), early stopping monitoring AUC on the validation set with a patience of five epochs, and best weights restored at the end of training.

### Data

DeepFisFis was trained and evaluated on 22 annotated recordings of C57BL/6 mice drawn from published and unpublished sources (Table 2). At 250 kHz, 19 recordings were used: eight pre-annotated recordings from AMVOC (Stoumpou et al., 2023), ten from USVSEG (Tachibana et al., 2020), and one recording from the mouseTube database (Torquet et al., 2016) annotated by the authors. At 384 kHz, three recordings were used: one pre-annotated recording from AMVOC and two recordings acquired and annotated in our laboratory during male–female pair interactions, previously collected for a separate study, using a CM16/CMPA condenser ultrasound microphone and an UltraSoundGate 116Un recording interface (Avisoft Bioacoustics, Germany) at 384 kHz and 16-bit resolution

**Table 2.** c. Datasets used for development and evaluation. Durations and call counts are derived from the supplied recordings and annotations.

| Source | Sampling rate (kHz) | # Recordings | Duration | # USVs | Annotation |
| --- | --- | --- | --- | --- | --- |
| AMVOC (Stoumpou et al., 2023) | 250 (384) | 8 (1) | ~46 s (6 s) | 203 (42) | public |
| USVSEG (Tachibana et al., 2020) | 250 | 10 | ~19.9 min | 2,799 | public |
| mouseTube (Torquet et al., 2016) | 250 | 1 | 10 min | 291 | authors |
| Laboratory (unpublished) | 384 | 2 | ~93.1 min | 6,935 | authors |

For cross-validation, recordings at each sampling rate were separately concatenated, divided into consecutive 10 s parts, and assigned to five folds. The 250 kHz folds were used for both training and evaluation. The 384 kHz folds were used for evaluation only.

### Evaluation and comparison

We employed two complementary approaches to score detections. *Temporal* scoring compares predicted and annotated calls sample by sample along the recording and reports precision, recall, and F1 score from the per-sample overlap between predicted and annotated calls. *Event* scoring operates at the level of whole calls: we define a True Positive Event (TPE) as an annotated call that overlaps any predicted call, and a True Discovery (TD) as a predicted call that overlaps any annotated call. Unmatched annotated calls are False Negative Events (FNE) and unmatched predicted calls are False Discoveries (FD). We define recall as the fraction of annotated calls that were matched (TPE out of all annotated calls), precision as the fraction of predicted calls that were matched (TD out of all predicted calls), and F1 score as the harmonic mean of the two.

Processing speed was measured as the time to execute the full detection pipeline (‘Detection pipeline’), averaging over 1000 repetitions, at input lengths of 5 ms, 400 ms, and 10 s. Timing was performed on a laptop (Intel Core i7-12800H, NVIDIA RTX 3080 Ti Laptop GPU, 64 GB RAM).

For comparison, DeepSqueak detections (Coffey et al., 2019) were scored with the same two schemes. Detections were obtained for each recording either from those distributed with the source dataset or, where these were unavailable, by running DeepSqueak v3.1 in MATLAB R2021b with the supplied Mouse Detector YOLO R2 network and default detection settings. Processing speed was also compared at these input lengths. DeepSqueak was not applicable at 5 ms, since the shortest input it accepted in our testing was 400 ms, and was therefore compared at 400 ms and 10 s only.

### Closed-loop implementation and latency measurement

Closed-loop operation was tested in a playback configuration. Previously recorded USVs were played through an ultrasonic electrostatic speaker (ESS16, Avisoft Bioacoustics, Germany) driven by an UltraSoundGate Player 216H, re-recorded with a Pettersson M500-384 USB ultrasound microphone (Pettersson Elektronik AB, Sweden) sampling at 384 kHz with 16-bit resolution, and processed by DeepFisFis using one of the five fine-tuned 384 kHz cross-validation models, in a continuous stream of 10 ms chunks (3,840 samples) acquired with PyAudio through the Windows WASAPI host API, a chunk duration bound by the microphone and host audio API used here. One positively classified segment was sufficient to trigger immediate playback of an 18 kHz tone through the internal speaker of the same laptop, chosen below the ultrasonic range so that it could not interfere with detection. Both the played-back vocalizations and the triggered tone were captured by the same microphone, so that their relative timing was preserved in a single recording. System latency was defined as the delays imposed by the hardware and software components of the setup, independently of the detector. These comprise audio chunk duration, microphone latency, software overhead, speaker latency and acoustic propagation delay. It was measured in the same configuration with DeepFisFis bypassed, by triggering the tone on arrival of the first audio chunk (Figure 3A), averaged over 1000 repetitions.

### Event-triggered recording analysis

Event-triggered recording was evaluated offline on the 384 kHz recordings using the detections produced by the 384 kHz model. For a given buffer size, each detected event was expanded by that buffer on both sides, overlapping intervals were merged, and two quantities were computed: the retained recording time as a fraction of total recording time, and the annotated USV time falling inside retained intervals as a fraction of total annotated USV time. Buffer sizes were varied logarithmically from 5 ms to 10 s and the analysis was repeated for each cross-validation fold (Figure 4B).

### Implementation and statistics

All training, evaluation and analysis were performed with custom Python 3.10 scripts, using TensorFlow/Keras 2.10 (Abadi et al., 2016) for model implementation and training, SciPy (Virtanen et al., 2020) and librosa (McFee et al., 2015) for signal processing, NumPy (Harris et al., 2020) and pandas (McKinney, 2010) for data handling, scikit-learn (Pedregosa et al., 2011) for cross-validation splitting, and Matplotlib (Hunter, 2007) and seaborn (Waskom, 2021) for figure generation. No inferential statistical tests were performed: all reported values are descriptive summaries (mean ± SD) across the five cross-validation folds for detection metrics, or across repetitions for timing and latency measurements, and the number of folds or repetitions is stated in each figure legend. Versions of the remaining packages are specified in the environment file distributed with the analysis code (see Data availability).

### Generative AI assistance

Claude (Anthropic) and ChatGPT (OpenAI) were used, under author supervision, to edit manuscript text and to check internal consistency. They were not used to generate or analyse data, produce figures, or draw scientific conclusions. The authors reviewed all text and take full responsibility for the manuscript.

## Acknowledgements

This work was supported by German Research Foundation (SPP 2041 SCHW1578/2-1 and SPP 2411 SCHW 2100/1-1) to M.K.S.. Research in the Schwarz laboratory was also supported by the Verein zur Förderung der Epilepsieforschung e.V. (M.K.S.) and from the program Netzwerke 2021 (iBehave, grant no. NW21-049), an initiative of the Ministry of Culture and Science of the State of North Rhine-Westphalia (M.K.S.).

We thank Frank Kurth for supervising the MSc thesis of A.M. at the University of Bonn, Institute of Computer Science.

## Author contributions

Ali Mohammadi (AM), Jens Tillmann (JT), Martin K. Schwarz (MKS)

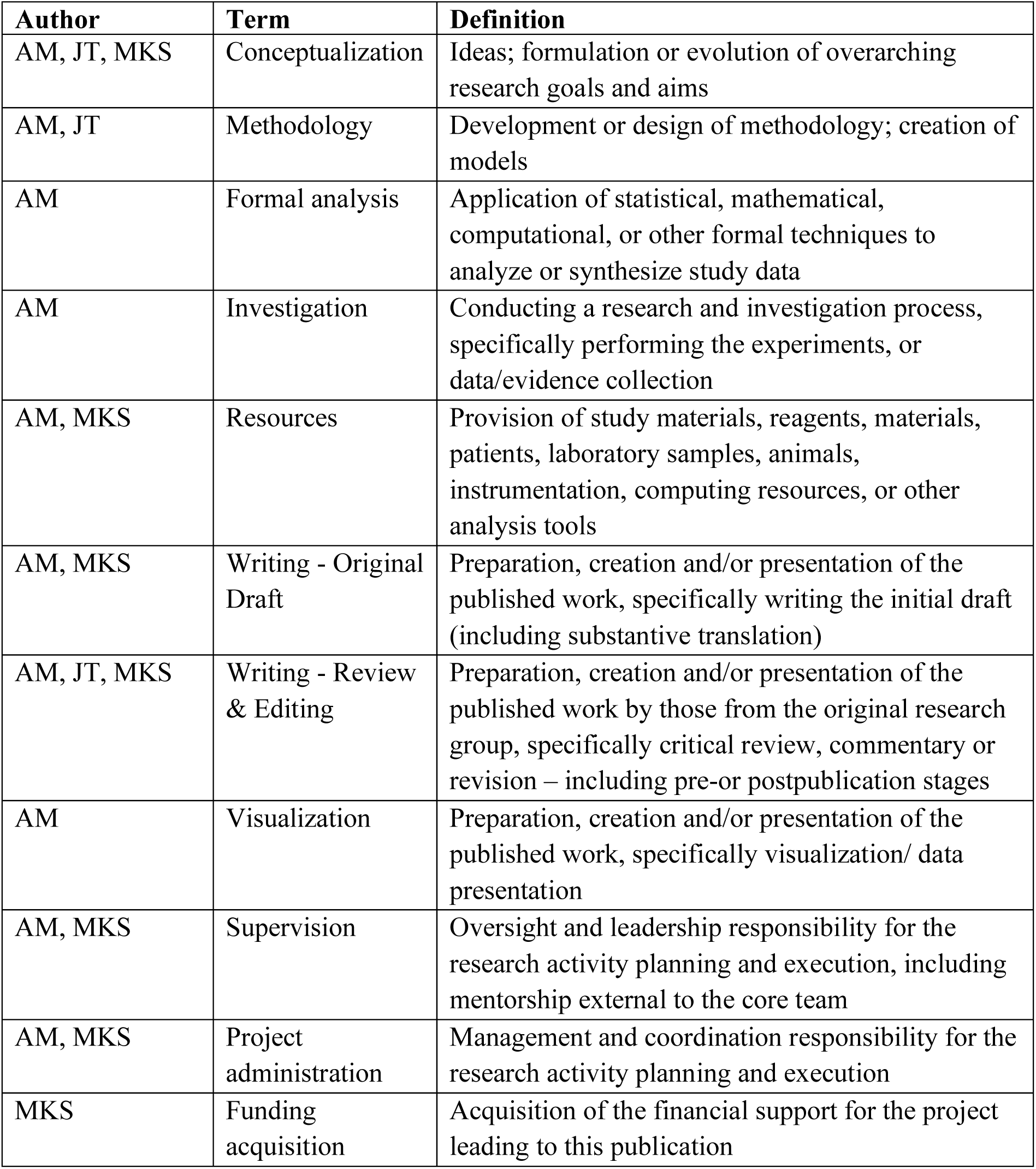

| Author | Term | Definition |
| --- | --- | --- |
| AM, JT, MKS | Conceptualization | Ideas; formulation or evolution of overarching research goals and aims |
| AM, JT | Methodology | Development or design of methodology; creation of models |
| AM | Formal analysis | Application of statistical, mathematical, computational, or other formal techniques to analyze or synthesize study data |
| AM | Investigation | Conducting a research and investigation process, specifically performing the experiments, or data/evidence collection |
| AM, MKS | Resources | Provision of study materials, reagents, materials, patients, laboratory samples, animals, instrumentation, computing resources, or other analysis tools |
| AM, MKS | Writing - Original Draft | Preparation, creation and/or presentation of the published work, specifically writing the initial draft (including substantive translation) |
| AM, JT, MKS | Writing - Review & Editing | Preparation, creation and/or presentation of the published work by those from the original research group, specifically critical review, commentary or revision – including pre-or postpublication stages |
| AM | Visualization | Preparation, creation and/or presentation of the published work, specifically visualization/ data presentation |
| AM, MKS | Supervision | Oversight and leadership responsibility for the research activity planning and execution, including mentorship external to the core team |
| AM, MKS | Project administration | Management and coordination responsibility for the research activity planning and execution |
| MKS | Funding acquisition | Acquisition of the financial support for the project leading to this publication |

## Ethics

Publicly available recordings from AMVOC (Stoumpou et al., 2023), USVSEG (Tachibana et al., 2020) and mouseTube (Torquet et al., 2016) were re-analysed. The 384 kHz recordings analysed here were acquired previously, as part of a larger dataset collected for a separate study, under animal licence AZ 81-02.04.2022.A137 approved by Landesamt für Natur, Umwelt und Verbraucherschutz Nordrhein-Westfalen.

## Data availability

AMVOC (Stoumpou et al., 2023) and USVSEG (Tachibana et al., 2020) are publicly available. The mouseTube recording (file C57_pair1_2100.wav) is available from the mouseTube database (Torquet et al., 2016) at https://mousetube.pasteur.fr/. The DeepFisFis source code, trained models, and the scripts used to generate all figures will be made publicly available on GitHub, with an archived release on bonndata, upon publication in a peer-reviewed journal. Until then, they are available from the corresponding author upon request.

